# Fast Non-Parametric Estimation of Outbreak Growth from Cumulative Incidence Applied to the Current Ebola Epidemic

**DOI:** 10.1101/340067

**Authors:** Thomas House

**Affiliations:** School of Mathematics, University of Manchester, M13 9PL, UK.

## Abstract

A method is presented that works with cumulative incidence data to provide a real-time estimate of the growth rate of an outbreak, without assuming any particular disease dynamics, and this is applied to the current Ebola outbreak.

## Working with Cumulative Incidence Data

During an ongoing outbreak, data are often not available at the level of detail that would be ideal, and in fact often the only publicly available data is on cumulative incidence – i.e. the times that new cases became symptomatic, but not recovery or infection times.

Previous work on Ebola has in fact required much more data than cumulative incidence to perform useful modelling [7, 5, 2, 1], however the question is then posed as to what can be done with cumulative incidence data while bearing in mind the limitations of naive approaches [4].

In this paper I outline an approach to cumulative incidence data that is part of a general framework I am developing called Time-Asymmetric Conjugate Statistical (TACS) learning. One paper has been submitted [3] on a specific, Bernoulli, case of this procedure. A more comprehensive manuscript applying the approach to many different datasets and providing careful comparisons with other methods is in preparation, however since this method may be useful to an unfolding public-health crisis I am releasing full mathematical details and code for the approach, together with results on the current Ebola outbreak, with the caveat that the work is somewhat preliminary.

The TACS estimation approach outlined here rests on three main assumptions:

1. The Force of Infection follows a Gamma distribution
2. Bayes’ theorem is a good update rule given new information
3. These two ingredients are all that is required

Of these, 1 and 2 are the strongest – the Gamma distribution is quite flexible, and Bayes’ theorem is sound – but 3 is weak since ideally we would be able to build in more scientific knowledge. As such, the current approach is in no way a substitute for a full transmission-dynamic model, but is rather a way to ‘make the best of’ limited data availability.

Despite this, applying the method to data on the current outbreak gives the results below, with the most important being Figure 4, which shows close to zero current and past growth, but very fast growth around two weeks in to the outbreak. Roughly speaking, a significant trend above the red line implies growth, and a significant trend below implies control.

## Mathematical results

The Gamma distribution has pdf

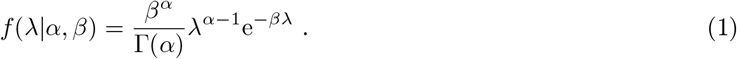

The probability of observing *y* new cases over time *δt* if the force of infection is λ is Poisson:

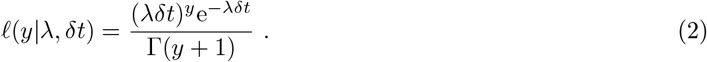

Bayes’ theorem holds for any conditional densities and states

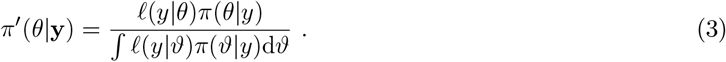

Applying this to our Gamma prior and Poisson likelihood gives that

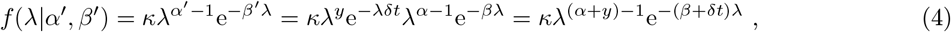

where *κ* is a normalising constant independent of λ, which gives us the update rules

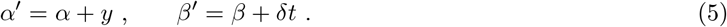

If λ is Gamma distributed with time-dependent parameters *α*(*t*) and *β*(*t*), then *X* = ln(λ) has pdf

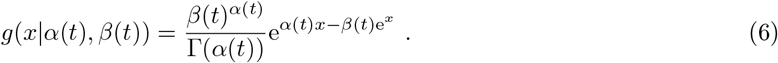

Integrating gives

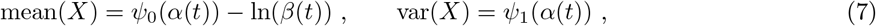

where *ψ_m_* is the polygamma function of order *m*. If we consider the effective growth rate *r* to be the derivative of *X*, then we obtain its mean and variance as

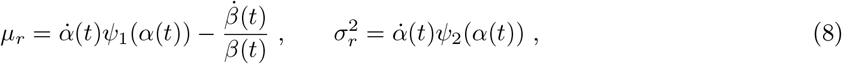

where a dot stands for differentiation with respect to time. These can be estimated at observation points as follows.

We assume that our data is composed of sequential observations of cumulative incidence *C*_1_, *C*_2_,…, *C_n_* at time points *t*_1_,*t*_2_,…*t_n_* such that *t_i_* < *t*_*i*+1_, ∀*i*. We then write our update rules as functions of time:

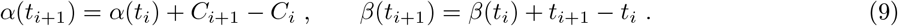

This leaves open the values *α*(0), *β*(0). This can be done through maximum likelihood. Firstly we note that

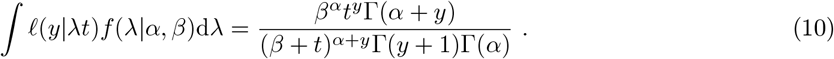

Then letting *t*_0_ = 0, *C*_0_ = 0 we have likelihood

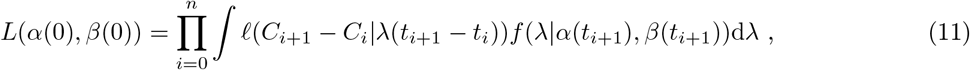

and let

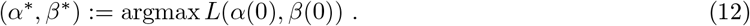

We numerically optimise *L*, then run the update rules at the optimal initial conditions, before estimating derivaties of the moments of the growth rate *r*.

**Fig 1:**
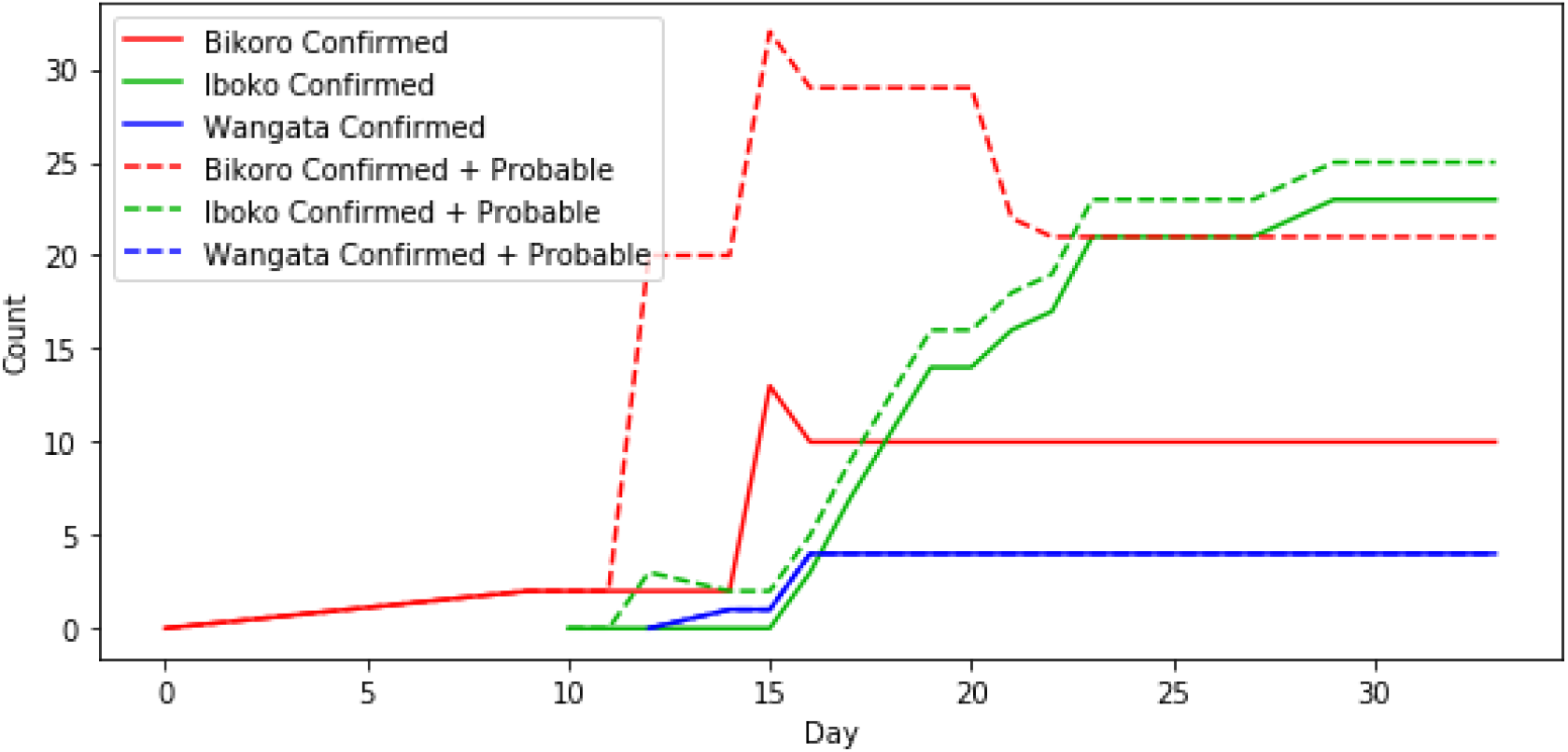
Cases. Confirmed and Probable Cases [6].

**Fig 2:**
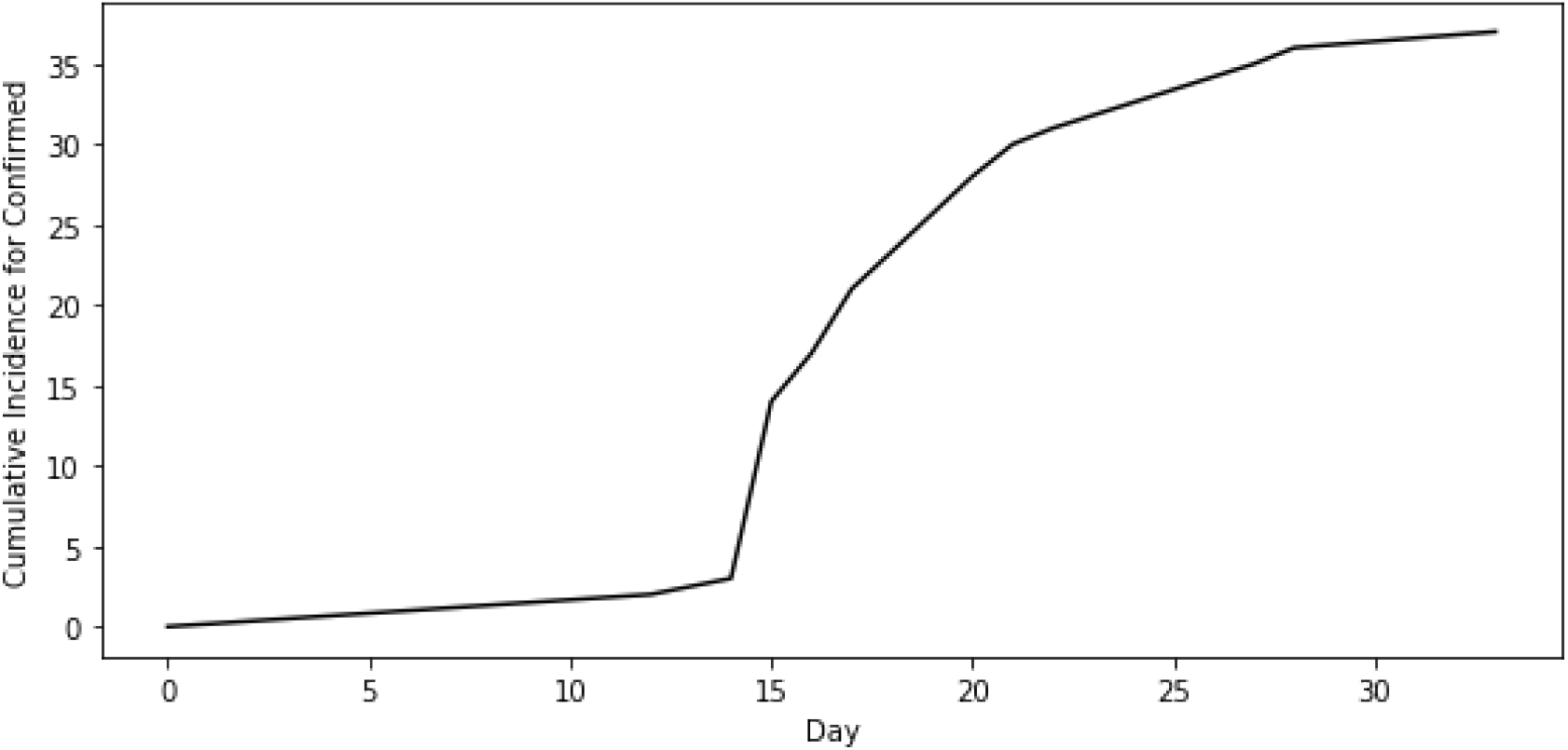
Cumulative Incidence. Assumed cumulative incidence curve.

**Fig 3:**
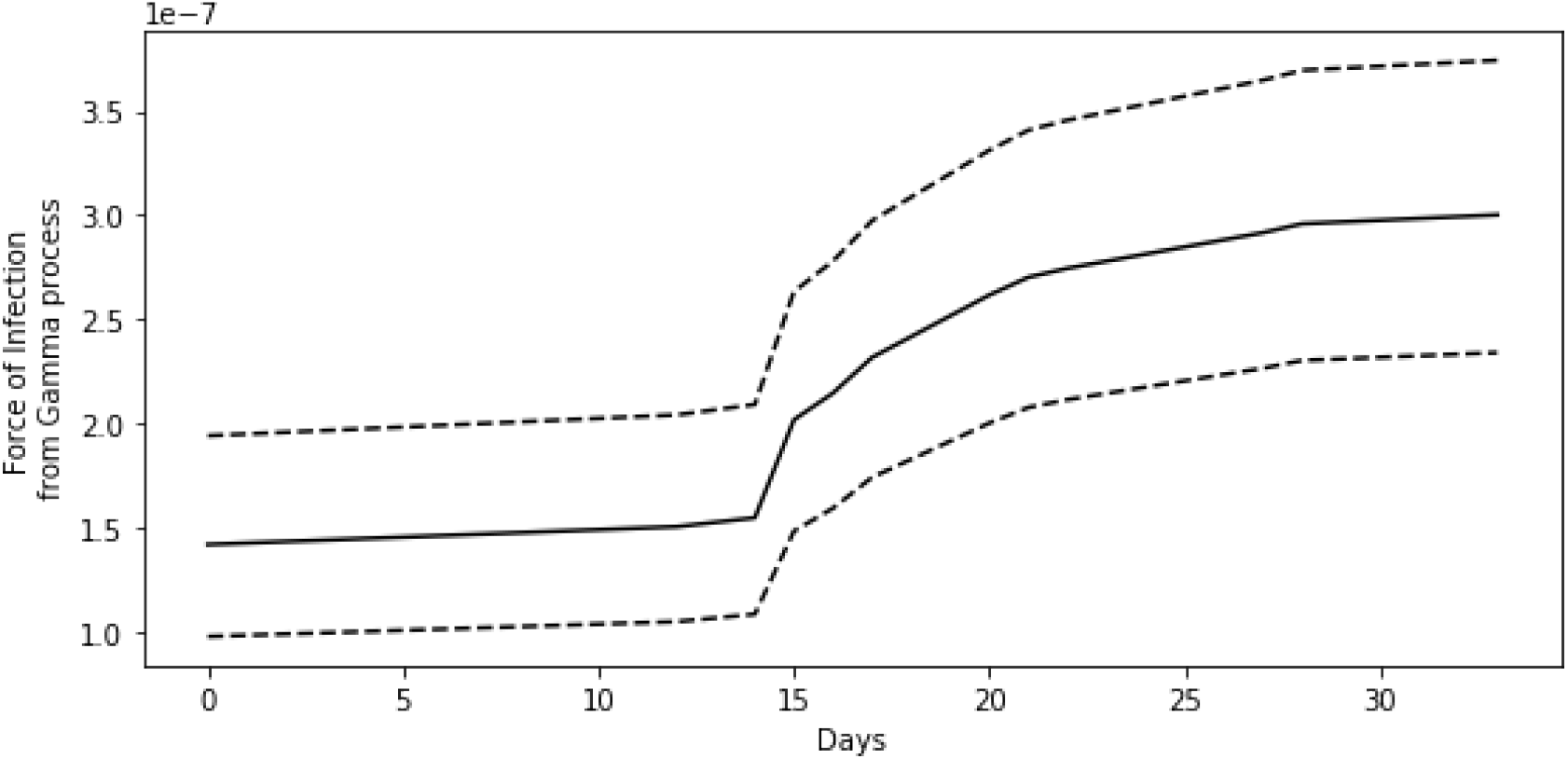
FOI. Inferred Force of Infection, λ.

**Fig 4:**
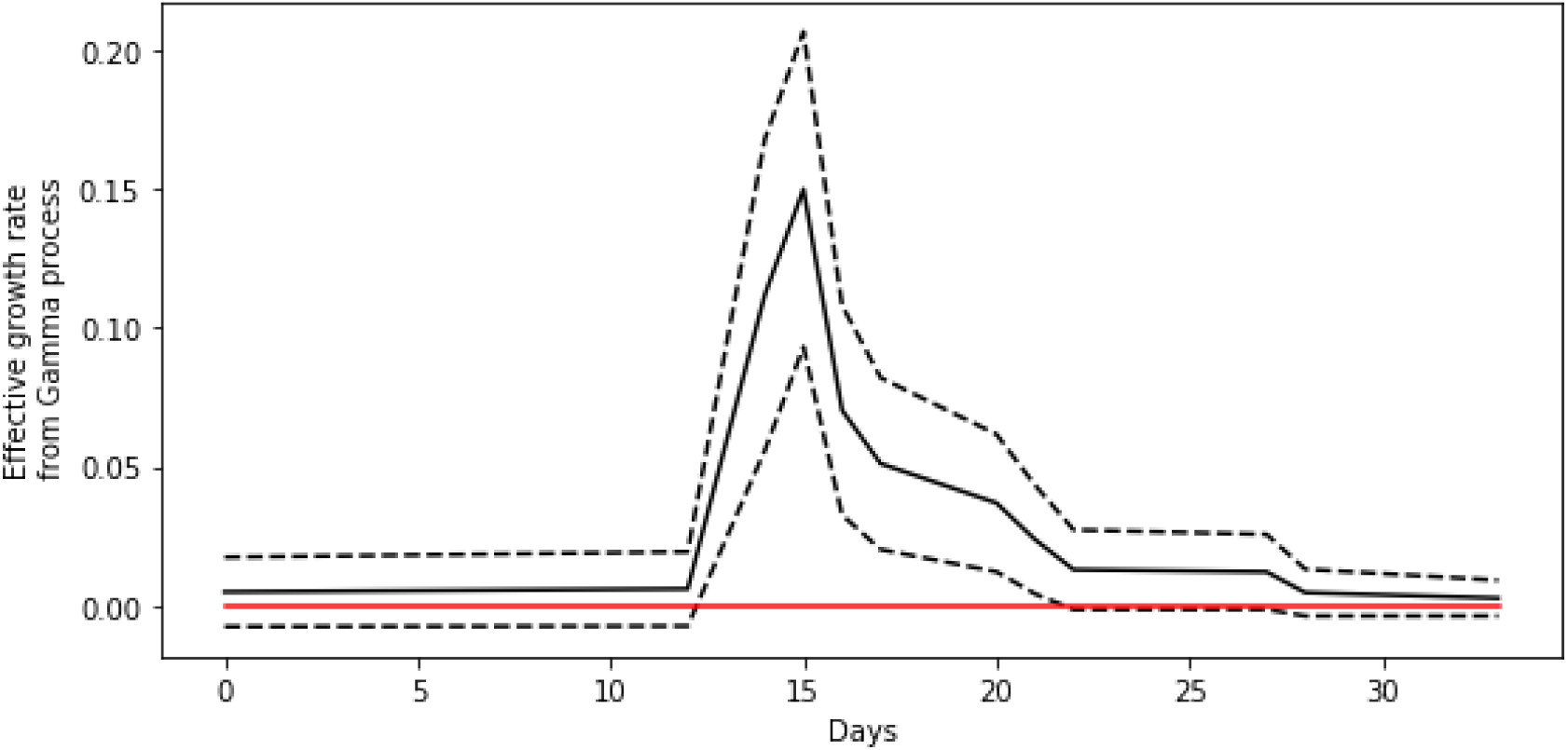
Growth. Inferred growth rate *r*.

## Funding Statement

Work supported by the Engineering and Physical Sciences Research Council, Grant Number EP/N033701/1.

## Python Code and Results

~~~
In [1]: %matplotlib inline
           import numpy as np
           import scipy.stats as st
           import scipy.special as sp
           import csv
           import matplotlib.pyplot as plt
           import urllib
           from io import StringIO
           from datetime import datetime
           import scipy.optimize as op
~~~

~~~
In [2]: url = ‘https://raw.githubusercontent.com/cmrivers/ebola_drc/master/drc/data.csv’
        uo = urllib.request.urlopen(url)
        ub = uo.read()
        us = ub.decode().replace(“\r\n”, “\n”)
        print(us)
~~~

~~~
event_date,report_date,health_zone,confirmed_cases,new_confirmed,probable_cases,new_probable,suspect_cases,new_suspect,deaths
20180501.20180510,Bikoro,,,,,21,,17
20180510.20180510,Bikoro,2,,,,9,,1
20180511.20180511,Bikoro,2,,,,6,,0
20180511,20180511,Iboko,0,,,,6,,1
20180512.20180512,Bikoro,2,,,,6,,0
20180512,20180512,Iboko,0,,,,6,,1
20180513.20180514,Bikoro,2,0,18,0,12,0,
20180513,20180514,Iboko,0,0,3,0,5,2,
20180513.20180514,Wangata,0,0,0,0,2,0,
20180515.20180516,Bikoro,2,0,18,0,15,1,
20180515,20180516,Iboko,0,0,2,0,3,0,
20180515.20180516,Wangata,1,0,0,0,3,0,
20180515.20180516,Ntondo,0,0,0,0,0,0,
20180516.20180517,Bikoro,13,0,19,1,4,0,21
20180516,20180517,Iboko,0,0,2,0,3,0,3
20180516.20180517,Wangata,1,0,0,0,3,0,1
20180516.20180517,Ntondo,0,0,0,0,0,0,0
20180517.20180518,Bikoro,10,0,19,0,0,0,21
20180517,20180518,Iboko,3,0,2,0,4,4,3
20180517.20180518,Wangata,4,0,0,0,1,1,1
20180518.20180520,Bikoro,10,0,19,0,0,0,22
20180518,20180520,Iboko,7,0,2,0,2,2,3
20180518.20180520,Wangata,4,0,0,0,2,1,1
20180520.20180521,Bikoro,10,0,19,0,0,0,22
20180520,20180521,Iboko,14,0,2,0,0,0,3
20180520.20180521,Wangata,4,0,0,0,2,2,2
20180521.20180522,Bikoro,10,0,19,0,0,0,23
20180521,20180522,Iboko,14,0,2,0,6,6,3
20180521.20180522,Wangata,4,0,0,0,3,2,1
20180522.20180523,Bikoro,10,0,12,1,2,2,16
20180522,20180523,Iboko,16,2,2,0,6,4,3
20180522.20180523,Wangata,4,0,0,0,6,3,3
20180523.20180524,Bikoro,10,0,11,0,2,2,16
20180523,20180524,Iboko,17,1,2,0,5,1,3
20180523.20180524,Wangata,4,0,0,0,1,0,
20180524.20180525,Bikoro,10,0,11,0,3,3,16
20180524,20180525,Iboko,21,4,2,0,2,0,4
20180524.20180525,Wangata,4,0,0,0,1,1,3
20180525.20180526,Bikoro,10,0,11,0,2,2,16
20180525,20180526,Iboko,21,0,2,0,2,0,6
20180525.20180526,Wangata,4,0,0,0,1,1,3
20180525.20180526,Ntondo,0,0,0,0,1,1,0
20180526.20180527,Bikoro,10,0,11,0,1,1,16
20180526,20180527,Iboko,21,0,2,0,5,3,6
20180526.20180527,Wangata,4,0,0,0,1,0,3
20180526.20180527,Ntondo,0,0,0,0,1,0,0
20180527.20180528,Bikoro,10,0,11,0,1,0,16
20180527,20180528,Iboko,21,0,2,0,4,0,6
20180527.20180528,Wangata,4,0,0,0,1,1,3
20180527.20180528,Ntondo,0,0,0,0,0,0,0
20180528.20180529,Bikoro,10,0,11,0,1,0,16
20180528,20180529,Iboko,21,0,2,0,1,1,6
20180528.20180529,Wangata,4,0,0,0,1,1,3
20180529.20180530,Bikoro,10,0,11,0,0,0,16
20180529,20180530,Iboko,22,1,2,0,3,3,6
20180529.20180530,Wangata,4,0,0,0,1,0,3
20180530.20180531,Bikoro,10,0,11,0,0,0,17
20180530,20180531,Iboko,23,1,2,0,0,0,5
20180530.20180531,Wangata,4,0,0,0,0,0,3
20180531.20180601,Bikoro,10,0,11,0,3,3,17
20180531,20180601,Iboko,23,0,2,0,0,0,5
20180531.20180601,Wangata,4,0,0,0,2,2,3
20180601.20180602,Bikoro,10,0,11,0,2,2,17
20180601,20180602,Iboko,23,0,2,0,3,3,5
20180601.20180602,Wangata,4,0,0,0,2,2,3
20180602.20180603,Bikoro,10,0,11,0,0,0,17
20180602,20180603,Iboko,23,0,2,0,3,1,5
20180602.20180603,Wangata,4,0,0,0,0,0,3
20180603.20180604,Bikoro,10,0,11,0,5,5,17
20180603,20180604,Iboko,23,0,2,0,0,0,5
20180603.20180604,Wangata,4,0,0,0,1,1,3
~~~

~~~
In [3]: reader = csv.reader(Stringĩũ(us), delimiter=’,')
        next(reader, None) *#Header*
        tb = np.array([])
        ti = np.array([])
        tw = np.array([])
        ib = np.array([])
        ii = np.array([])
        iw = np.array([])
        ibp = np.array([])
        iip = np.array([])
        iwp = np.array([])
        for row in reader:
            if row[2] == ‘Bikoro’:
               tb = np.append(tb,datetime.strptime(row[0],‘%Y%m%d’).timestamp())
               ib = np.append(ib,row[3])
               ibp = np.append(ibp,row[5])
            elif row[2] == ‘Iboko’:
               ti = np.append(ti,datetime.strptime(row[0],‘%Y%m%d’).timestamp())
               ii = np.append(ii,row[3])
               iip = np.append(iip,row[5])
            elif row[2] == ‘Wangata’:
               tw = np.append(tw,datetime.strptime(row[0],‘%Y%m%d’).timestamp())
               iw = np.append(iw,row[3])
               iwp = np.append(iwp,row[5])

        ib[ib== ' '] = ' 0 '
        ii[ii=='']='0'
        iw[iw==''] = '0'
        ibp[ibp=='']='0'
        iip[iip=='']='0'
        iwp[iwp=='']='0'
        t0 = np.min(np.concatenate((tb,ti,tw)))
        tb = np.round((tb-t0)/(60*60*24))
        ti = np.round((ti-t0)/(60*60*24))
        tw = np.round((tw-t0)/(60*60*24))
        ib = ib.astype(np.float)
        ii = ii.astype(np.float)
        iw = iw.astype(np.float)
        ibp = ibp.astype(np.float)
        iip = iip.astype(np.float)
        iwp = iwp.astype(np.float)
~~~

~~~
In [14]: plt.figure(figsize=(8, 4))
         plt.plot(tb,ib,c=[1,0,0])
         plt.plot(ti,ii,c=[0,0.7,0])
         plt.plot(tw,iw,c=[0,0,1])
         plt.plot(tb,ib+ibp,c=[1,0,0],linestyle='dashed')
         plt.plot(ti,ii+iip,c=[0,0.7,0],linestyle='dashed')
         plt.plot(tw,iw+iwp,c=[0,0,1],linestyle='dashed')
         plt.legend(('Bikoro Confirmed','Iboko Confirmed','Wangata Confirmed',
                     'Bikoro Confirmed + Probable','Iboko Confirmed + Probable',
                     'Wangata Confirmed + Probable'))
         plt.xlabel('Day')
         plt.ylabel('Count')
         plt.tight_layout()
~~~

~~~
In [5]: *# Add confirmed cases*
           tt = np.union1d(tb,np.union1d(ti,tw))
           cc = 0*tt
           n = len(tt)
           for t in tb:
               j = np.where(tt==t)
               cc[j] += ib[tb==t]
           for t in ti:
               j = np.where(tt==t)
               cc[j] += ii[ti==t]
           for t in tw:
               j = np.where(tt==t)
               cc[j] += iw[tw==t]
~~~

~~~
In [6]: *# Remove non-informative points - this assumes no doubly flat lines*
           kk = np.where(np.diff(cc)>0)
           kk = np.append(kk,len(cc)-1)
           tt = tt[kk]
           cc = cc[kk]
           n = len(tt)
~~~

~~~
In [15]: *# Plot*
            plt.figure(figsize=(8, 4))
            plt.plot(tt,cc,c=[0,0,0])
            plt.xlabel('Day')
            plt.ylabel('Cumulative Incidence for Confirmed')
            plt.tight_layout()
~~~

~~~
In [8]: def nllfun(x,tt,cc,nn):
            t = np.zeros_like(cc)
            y = np.zeros_like(cc)
            al = np.zeros_like(cc)
            bt = np.zeros_like(cc)
            al[0] = x[0]
            bt[0] = x[1]
            nll=0
            for i in range(1,nn):
                y[i] = cc[i] - cc[i-1]
                t[i] = tt[i] - tt[i-1]
                al[i] = al[i-1]
                bt[i] = bt[i-1]
                al[i] += y[i]
                bt[i] += t[i]
                nll += y[i] *np. log(t[i] / (t[i] +bt[i])) - al[i]*np.log((bt[i]/(t[i]+bt[i])))
                nll += sp.gammaln(al[i]+y[i]) - sp.gammaln(1+y[i]) - sp.gammaln(al[i])
return nll
~~~

~~~
In [9]: *#Optimizer to fit the initial conditions*
           nll = lambda xx: nllfun(np.abs(xx),tt,cc,n)
           fout = op.minimize(nll,np.array((1.0,1.0)),method='Nelder-Mead')
           astar = np.abs(fout.x[0])
           bstar = np.abs(fout.x[1])
           fout.x
~~~

~~~
Out[9]: array([-3.32222615e+01, 2.33945606e+08])
In [10]: t = np.zeros_like(cc)
         y = np.zeros_like(cc)
         al = np.zeros_like(cc)
         bt = np.zeros_like(cc)
         lam = np.zeros_like(cc)
         lal = np.zeros_like(cc)
         lau = np.zeros_like(cc)
         al[0] = astar
         bt[0] = bstar
         lam[0] = st.gamma.mean(al[0],0,1/bt[0])
         lal[0] = st.gamma.ppf(0.025,al[0],0,1/bt[0])
         lau[0] = st.gamma.ppf(0.975,al[0],0,1/bt[0])
~~~

~~~
for i in range(1,n):
    y[i] = cc[i] - cc[i-1]
    t[i] = tt[i] - tt[i-1]
    al[i] = al[i-1]
    bt[i] = bt[i-1]
    al[i] += y[i]
    bt[i] += t[i]
    lam[i] = st.gamma.mean(al[i],0,1/bt[i])
    lal[i] = st.gamma.ppf(0.025,al[i],0,1/bt[i])
    lau[i] = st.gamma.ppf(0.975,al[i],0,1/bt[i])
~~~

~~~
In [16]: plt.figure(figsize=(8, 4))
         plt.plot(tt,lam,c=[0, 0, 0])
         plt.plot(tt,lal,c=[0, 0, 0],linestyle='dashed')
         plt.plot(tt,lau,c=[0, 0, 0],linestyle='dashed')
         plt.xlabel('Days')
         plt.ylabel('Force of Infection\n from Gamma process')
         plt.tight_layout()
~~~

~~~
In [12]: am = al
         bm = bt
         dt = np.gradient(tt)
         dth = (np.gradient(bm)/(-1.0*bm**2))/dt
         da = (np.gradient(am))/dt
         rm = (dth*bm) + (sp.polygamma(1,am)*da)
         rv = (sp.polygamma(2,am)*da)
         rs = np.sqrt(-rv)
~~~

~~~
In [17]: plt.figure(figsize=(8, 4))
         plt.plot(tt,rm,c=[0,0,0])
         plt.plot(tt,rm-rs,c=[0,0,0],linestyle='dashed')
         plt.plot(tt,rm+rs,c=[0,0,0],linestyle='dashed')
         plt.plot(tt,0*tt,c=[1,0,0])
         plt.xlabel('Days')
         plt.ylabel('Effective growth rate\n from Gamma process')
         plt.tight_layout()
~~~

